# Single-nucleus RNA-seq reveals dysregulation of striatal cell identity due to Huntington’s disease mutations

**DOI:** 10.1101/2020.07.08.192880

**Authors:** Sonia Malaiya, Marcia Cortes-Gutierrez, Brian R. Herb, Sydney R. Coffey, Samuel R.W. Legg, Jeffrey P. Cantle, Carlo Colantuoni, Jeffrey B. Carroll, Seth A. Ament

## Abstract

Huntington’s disease (HD) is a dominantly inherited neurodegenerative disorder caused by a trinucleotide expansion in exon 1 of the huntingtin (*Htt*) gene. Cell death in HD occurs primarily in striatal medium spiny neurons (MSNs), but the involvement of specific MSN subtypes and of other striatal cell types remains poorly understood. To gain insight into cell type-specific disease processes, we studied the nuclear transcriptomes of 4,524 cells from the striatum of a genetically precise knock-in mouse model of the HD mutation, *Htt^Q175/+^*, and from wildtype controls. We used 14-15-month-old mice, a time point roughly equivalent to an early stage of symptomatic human disease. Cell type distributions indicated selective loss of D2 MSNs and increased microglia in aged *Htt^Q175/+^* mice. Thousands of differentially expressed genes were distributed across most striatal cell types, including transcriptional changes in glial populations that are not apparent from RNA-seq of bulk tissue. Reconstruction of cell typespecific transcriptional networks revealed a striking pattern of bidirectional dysregulation for many cell type-specific genes. Typically, these genes were repressed in their primary cell type, yet de-repressed in other striatal cell types. Integration with existing epigenomic and transcriptomic data suggest that partial loss-of-function of the Polycomb Repressive Complex 2 (PRC2) may underlie many of these transcriptional changes, leading to deficits in the maintenance of cell identity across virtually all cell types in the adult striatum.

## INTRODUCTION

Huntington’s disease (HD) is a fatal neurodegenerative disorder caused by dominant inheritance of trinucleotide repeat expansion mutations in the huntingtin (*HTT*) gene^1^. Clinical symptoms include deficits in motor control and cognition, as well as psychiatric symptoms. Although the causal mutation has been known for >25 years, there are no existing treatments that dramatically alter disease progression. In the absence of treatment, symptoms progressively worsen, leading inevitably to death 10-15 years after the symptomatic age at onset.

An enduring mystery in HD biology is why Huntington’s disease mutations lead to selective neurodegeneration in specific subtypes of neurons, while other nearby cells remain largely spared, despite the fact that the *HTT* gene is robustly expressed in most or all cell types. HD progression is linked to the selective cell death of medium spiny neurons (MSNs) in the striatum^2^. Among MSNs, *Drd2-expressing* MSNs that project to the lateral segment of the globus pallidus (termed ‘D2 MSNs’) are thought to be more vulnerable than *Drd1*-expressing MSNs that project to the entopenduncular nucleus and the substantia nigra pars reticulata (termed ‘D1 MSNs’)^3^. Striatal interneurons are less vulnerable than MSNs but may undergo disease-related changes that remain little studied^4,5^. Neurodegeneration in HD is accompanied by neuroinflammatory processes. Microglial activation occurs prior to the clinical manifestation of the disease^6^. Reactive astrogliosis occurs relatively late in the natural progression of HD pathophysiology but may still contribute to neurotoxicity^7,8^. Changes in forebrain white matter and striatal oligodendrocytes begin early in disease progression, including deficits in myelinated axon fibers and increased oligodendrocyte cell number^9,10^. However, the regulation and impact of these inflammatory processes across glial and neuronal cell types remains poorly understood.

Transcriptional changes are among the earliest phenotypes in cells and tissues expressing mHTT and are highly reproducible in human HD^11–13^. MSN-specific genes and components of synapses are down-regulated, while up-regulated genes include signatures of neuroinflammation^7,14^. Notably, there is evidence that some transcriptional changes are directly related to functions of HTT in the nucleus, including interactions of both wildtype and mutant HTT with transcriptional regulatory proteins^15–17^. However, previous transcriptomic studies using bulk tissue failed to illuminate the cell type-specificity of many disease processes.

Single-cell transcriptomics has emerged as a scalable technology enabling an unprecedented view of cell types and cell states in the mammalian brain. To date, only a few published studies have applied this approach to any neurodegenerative disease^18–22^. Here, we analyzed the nuclear transcriptomes of 4,524 striatal cells from a genetically precise knock-in mouse model of a juvenile-onset HD mutation, *Htt^Q175/+^*. Our analyses of these data reveal numerous insights into cell type-specific disease processes.

## RESULTS

### Single-nucleus RNA-seq of 14-15 month-old Htt^Q175/+^ HD knock-in mice and wildtype controls

We generated single-nucleus RNA-seq from the striatum of four male 14-15 month-old *Htt^Q175/+^* mice and four male wildtype controls using the 10x Genomics Chromium system. *Htt^Q175/+^* is a widely used genetically precise mouse model for a mutation associated with juvenile-onset HD in which a humanized *HTT* exon 1 fragment with 140 CAG repeats was knocked into the endogenous *Htt* locus and the repeats spontaneously expanded to approximately 175 CAG repeats, which was later stabilized at ~190 repeats. These mice have normal lifespan, with progressive behavioral, neuroanatomical, and transcriptomic deficits^12,23,24^. At 14-15-months-old, striatal atrophy is detectable, but the presence of neuronal cell death has been controversial. This dataset represents, to our knowledge, the first single-nucleus RNA-seq study utilizing a genetically precise mouse model of the HD mutation. In addition, the mice in our study are considerably older than mice studied in previously published datasets from knock-in mouse models of the HD mutation, providing insights into a timepoint roughly equivalent to early manifest disease that has not been adequately modeled in previous studies utilizing these mouse models.

Following QC and normalization, we analyzed 4,524 high-quality cells, of which 3,210 were derived from *Htt^Q175/+^* mice and 1,314 from wildtype mice (Methods; Fig. S1). Louvain clustering and annotation with known marker genes^25^ revealed well-defined clusters corresponding to each of the major cell populations in the striatum, including 3,003 MSNs, 288 *Sst+* interneurons, 120 *Pvalb+* interneurons, 73 *Chat+* interneurons, 468 oligodendrocytes, 300 astrocytes, 112 endothelial cells, 82 microglia, and 78 polydendrocytes (Fig. 1a,b; Fig. S1; Table S1). Sub-clustering of MSNs using 36 sub-type marker genes with >8-fold differences in expression in prior scRNAseq of mouse striatum^25^ revealed 1,809 D1 MSNs and 941 D2 MSNs (Fig. 1c), as well as 166 MSNs whose expression profiles match the recently described “eccentric” subtype^25^. We have created a web portal for visualization and analysis of these data at the Gene Expression Analysis Resource (https://umgear.org/p?l=1d76bf3e).

**Figure 1.**
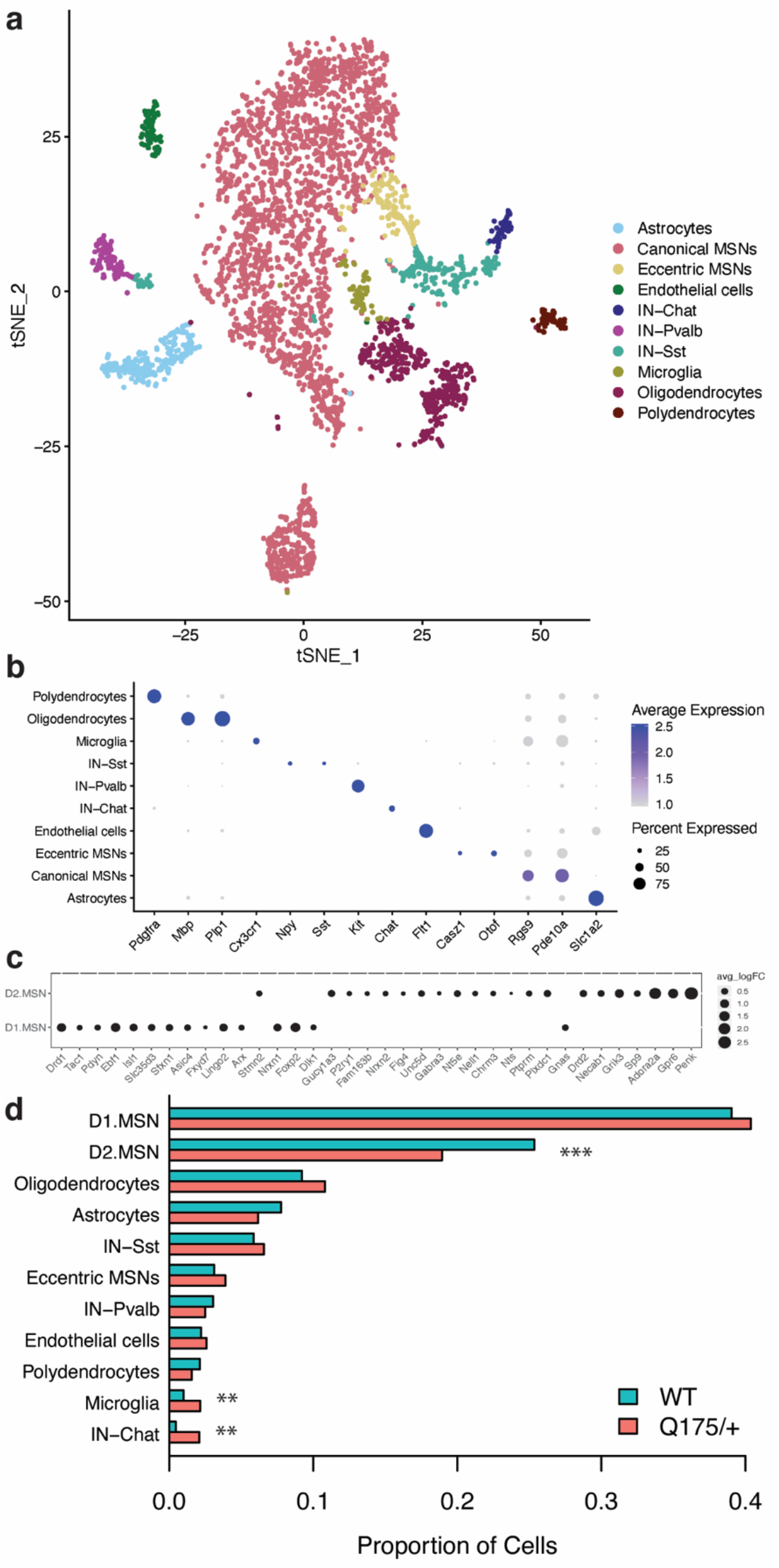
Identification of cell types in single-nucleus RNA-seq of 4,524 cells from the striatum of 14-15-month-old *Htt^Q175/+^* vs. *Htt^+/+^* mice. a. tSNE plot of major cell types. B. Dot plot of top marker genes for all cell types. c. Dot plot showing marker genes for D1 vs. D2 MSN sub-types. d. Proportion of cells from *Htt^Q175/+^* vs. *Htt^+/+^* mice in each cell type. Fisher’s exact test: *** p < 0.001; ** p < 0.01.

While most cell types were represented at similar proportions in *Htt^Q175/+^* vs. wildtype mice, several differences were noted (Fig. 1d). The proportion of cells identified as D2 MSNs was ~30% lower in *Htt^Q175/+^* mice (odds ratio = 0.69; p = 2.3e-6). This decreased proportion of D2 MSNs was robust across a range of QC and clustering parameters (Fig. S2). These data suggest that D2 but not D1 MSNs may die in the striatum of *HTT^Q175^* mice aged over one year. These results are consistent with the progression of MSN cell death in human HD^3^. However, previous studies had failed to detect cell death at earlier time points in knock-in mouse models of the HD mutation, a source of concern from a disease modeling perspective. We also observed significant increases (p < 0.01) in the proportion of microglia and Chat+ interneurons. The former may indicate microglial proliferation, while the significance of the latter is unknown.

### Cell type-specific gene expression changes in HD knock-in mice

Next, we studied celltype-specific gene expression changes in *Htt^Q175/+^* vs. wildtype mice. We identified 13,897 celltype-specific gene expression changes, involving 8,124 distinct genes (differentially expressed genes, DEGs; False Discovery Rate [FDR] < 0.05; Fig. 2a; Table S1). In our primary analysis, we detected DEGs by applying Wilcoxon rank-sum tests to smoothed read counts. A second approach, applying Wilcoxon signed-rank tests to non-smoothed read counts yielded a similar rank-ordering of DEGs but with reduced statistical power. Microglia and *Chat+* interneurons were excluded from this analysis due to insufficient cell numbers.

**Figure 2.**
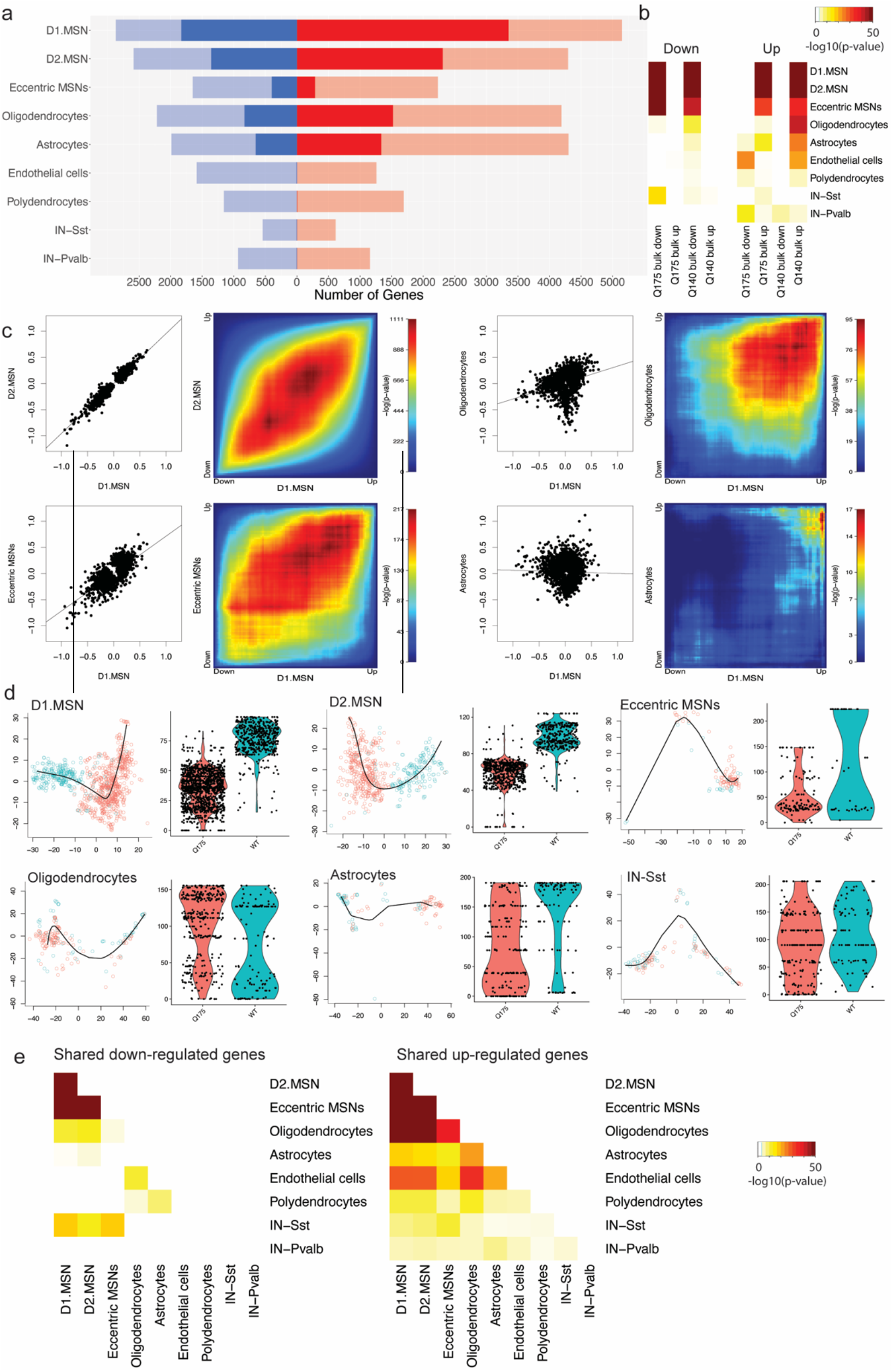
Characterization of differentially expressed genes in nine cell types from the striatum of 14-15-month-old *Htt^Q175/+^* vs. *Htt^+/+^* mice. a. Counts of differentially expressed genes in each cell type (red = up-regulated; blue = down-regulated; saturated color, FDR < 0.05; desaturated color, p < 0.01. b. Statistical overlap of up- and down-regulated genes with published RNA-seq of bulk striatal tissue from ten-month-old *Htt^Q175/+^* vs. *Htt^G20/+^* mice and from *Htt^Q140/+^* vs. *Htt^G20/+^* mice. c. Scatterplots and rank-rank hypergeometric overlap heatmaps indicating shared and unique gene expression changes in selected pairs of cell types. d. Slingshot pseudotime trajectory analysis in MSN subtypes, astrocytes, oligodendroocytes, and SST interneurons. e. Statistical overlap of up- and down-regulated genes among pairs of cell types.

We found 5,181, 3,666, and 685 DEGs in D1, D2, and eccentric MSNs, respectively. Comparison of these celltype-specific DEGs to previously described lists of DEGs from RNA-seq of bulk striatal tissue from ten-month-old *Htt^Q175/+^* mice vs. *Htt^Q20/+^* controls^12^ indicated that both up- and down-regulated DEGs in all MSN subtypes were strongly enriched for known DEGs from bulk tissue RNA-seq (limma geneSetTest: p < 1e-100 for DEGs in D1 and D2 MSNs; p < 1e-30 in eccentric MSNs; Fig. 2b; Table S3). Genes with the lowest p-values included *Pde10a, Rgs9, Wnt8b, Trank1, Scn4b, Rap1gap, Pde1b, Ptpn5, Adcy5, Atp2b1*, and *Arpp21*, all of which are also among the strongest and most-consistently observed DEGs from previous studies in bulk tissue. Down-regulated genes in MSNs were enriched for synaptic functions (e.g., “neuron spine”, p-values = 7.95e-7, 1.95e-8, and 8.1e-4 in D1, D2, and eccentric MSNs, respectively; Table S3). Up-regulated genes in MSNs were enriched for genes localized to the “nucleus” (p = 4.1e-11, 2.0e-10, and 0.014 in D1, D2, and eccentric MSNs, respectively), especially genes related to “histone modification” (p = 3.8e-4, 8.8e-4, and 3.1e-3 in D1, D2, and eccentric MSNs, respectively). Comparing the fold changes of DEGs in D1, D2, and eccentric MSNs revealed that with very few exceptions these fold changes were nearly identical in magnitude and direction (Fig. 2c, left; Pearson correlation comparing the log2(fold changes) of the top 10% of genes ranked by p-value in D1 vs. D2 MSNs, r = 0.97, p << 1e=308; D1 vs. eccentric MSNs, r = 0.77, p = 3.0e-242). The slope of the regression line is ~1 in all of these comparisons among MSN subtypes. Down-sampling analyses suggested that the somewhat weaker correlation coefficient in D1 vs. eccentric MSNs was primarily due to the smaller eccentric MSN sample size and is not biologically meaningful. Pseudotime trajectory analysis with Slingshot^26^ indicated that nearly all MSNs in these 14-15-month-old *Htt^Q175/+^* mice exist in a disease-specific transcriptional state that is never observed in wildtype mice (Fig 2d). Thus, single-nucleus RNA-seq revealed strong transcriptional effects of the HD mutation in all three MSN subtypes, including eccentric MSNs, yet the enhanced vulnerability of D2 vs. D1 MSNs is not reflected in the magnitude of transcriptional effects, at least not in the current dataset.

We found 2,351 DEGs in oligodendrocytes from *Htt^Q175/+^* vs. *Htt^+/+^ mice*. Neither up-nor down-regulated DEGs in oligodendrocytes strongly overlapped known DEGs from bulk tissue RNA-seq of 10-month-old *Htt^Q175/+^* vs. *Htt^Q20/+^* mice (p = 0.01, 0.06, respectively). By contrast, our DEGs in oligodendrocytes strongly overlapped DEGs from RNA-seq of sorted oligodendrocytes in the striatum of BACHD mice -- a transgenic mouse model of the HD mutation -- compared to wildtype controls^27^ (p = 6.6e-19, 1.8e-13, for down- and up-regulated DEGs, respectively). Down-regulated DEGs in oligodendrocytes were strongly enriched for oligodendrocyte-specific functions such as “myelin sheath” (p = 6.75e-14), as well as more basic cellular processes such as “structural constituent of ribosome” (p = 8.64e-12) and “mitochondrion” (p = 3.16e-10). Up-regulated DEGs were enriched for several categories of genes that are typically associated with neurons, including “ion channel complex” (p = 2.0e-8) and “synaptic membrane” (p = 1.2e-6). Trajectory analysis of oligodendrocytes suggested that oligodendrocytes exist on a continuum from normal to disease-associated states; i.e., in contrast to the discrete disease-associated transcriptional states of MSNs, the disease-associated transcriptional states in oligodendrocytes are also present in *Htt^+/+^* mice, but at a lower frequency (Fig. 2d). In summary, we identified thousands of reproducible DEGs in oligodendrocytes that are obscured in bulk tissue RNA-seq.

We found 1,987 DEGs in astrocytes from *Htt^Q175/+^* vs. *Htt^+/+^*mice. The up-regulated genes in astrocytes overlapped known up-regulated DEGs from bulk tissue RNA-seq of 10-month-old *Htt^Q175/+^* vs. *Htt^Q20/+^* mice (p = 2.3e-12), whereas down-regulated gene sets in astrocytes did not significantly overlap these known DEGs (p > 0.05). Prior work has revealed neurotoxic reactive astrocytes in post-mortem striatal brain tissue from HD patients, but their presence in mouse models of the HD mutation is controversial^7,8^. To identify reactive astrocytes in our dataset, we sub-clustered astrocytes on the basis of 87 genes previously shown to be induced in reactive astrocytes^28^, revealing a cluster of 134 reactive astrocytes, enriched for pan-reactive markers such as *Gfap* (p = 3.6e-14) and *Vim* (p = 1.2e-9), as well as two clusters of non-reactive astrocytes that do not express these markers (Fig. S2). While reactive astrocytes were present in both genotypes, they were significantly more abundant in *Htt^Q175/+^* mice (Fisher’s exact test: OR = 2.3; p = 0.001). Many reactive astrocytes expressed markers of the ‘A1’ neurotoxic sub-type (e.g., *H2-D1*, p = 1.2e-8), whereas very few cells expressed markers of ‘A2’ neuroprotective reactive astrocytes. While these results support the presence of neurotoxic reactive astrocytes in the striatum of *Htt^Q175/+^* mice aged >1 year, several analyses suggest that prototypical reactive astrogliosis explains only a subset of the HD-related transcriptional changes in astrocytes. Trajectory analyses showed a strong shift in astrocyte cell states in *Htt^Q175/+^* vs. *Htt^+/+^*mice (Fig. 2d), but this trajectory was only weakly correlated with reactive vs. non-reactive subtypes (Fig. S3). Instead, up-regulated DEGs in astrocytes were most strongly enriched for the GO term “synapse” (p = 8.1e-32), while down-regulated DEGs were most strongly enriched for GO terms related to transcriptional regulation (e.g., “negative regulation of transcription by RNA polymerase II”, p = 2.7e-9). Thus, there are profound changes in the transcriptomes of astrocytes from *Htt^Q175/+^* vs. *Htt^+/+^* mice, only some of which reflect known neuroinflammatory processes.

At an FDR < 0.05, we detected fewer than ten DEGs in Sst+ and Pvalb+ interneurons, endothelial cells, and polydendrocytes (Fig. 2a). Moreover, trajectory analysis indicated that the principal curve in these cell types was not correlated with genotype (p > 0.05). We note that although these cell types are relatively rare, we were able to detect hundreds of DEGs in comparably rare eccentric MSNs. Therefore, our data indicate that these cell types are less vulnerable to the transcriptional effects of the HD mutation.

Comparisons of gene expression changes across cell types detected a marked difference in cell type-specificity of down-regulated vs. up-regulated genes. Down-regulated genes were largely nonoverlapping across cell types, with the only strong overlaps occurring among MSN subtypes (Fig. 2e). Many of these down-regulated DEGs were ‘cell identity’ genes that are expressed specifically in that same cell type. That is, top genes down-regulated in MSNs included MSN marker genes such as *Ppp1r1b, Pde10a*, and *Rgs9*. Genes down-regulated in astrocytes were enriched for astrocyte marker genes (astrocyte marker genes (p-value = 1.11e-212), including *Hes5, Gjb6*, and *Ddhd1*. And genes down-regulated in oligodendrocytes were enriched for oligodendrocyte-specific genes (p-value = 3.32e-155), including *Mog, Gjb1*, and *Cldn11*.

By contrast, many up-regulated DEGs were shared across cell types. with statistically significant overlap among up-regulated DEGs in D1 MSNs, D2 MSNs, eccentric MSNs, astrocytes, oligodendrocytes, endothelial cells, polydendrocytes, and *Sst+* interneurons (Fig. 2c). 30 genes were up-regulated (FDR < 0.05) in five distinct cell types. These included inflammation-related genes such as colony stimulating factor 2 receptor subunit alpha (*Csf2ra*), histocompatibility 2, D region locus 1 (*H2-D1*), and myocardial infarction associated transcript (*Miat*), suggesting that some shared changes are due to the broadly acting effects of pro-inflammatory molecules such as cytokines. However, broadly up-regulated genes also included genes that are not typically associated with inflammation, including synaptic genes like the GABA_A_ receptor alpha 1 subunit (*Gabra1*) and the voltage-gated sodium channel alpha 9 subunit (*Scn9a*). Thus, up-regulation of certain transcripts across multiple striatal cell types is a prominent feature of gene expression changes in *Htt^Q175/+^* vs. *Htt^+/+^*mice, involving both neuroinflammation-related and non-neuroinflammation-related genes.

### Network analyses reveal principles of transcriptional dysregulation

We reconstructed and analyzed gene co-expression networks to gain deeper insight into the processes driving transcriptional dysregulation within and across cell types. Gene co-expression networks are widely employed in RNA-seq with bulk tissue, but standard methods such as WGCNA do not work well with scRNAseq, as the sparseness of the data masks gene-gene correlation structure^29^. To overcome this issue, we used knn-smoothing^30^ to impute read counts across cells (k=15, nPCs = 30). We confirmed that this approach produced strong correlations among known markers of D1 MSNs and among known markers of D2 MSNs, without inducing spurious correlations among markers across subtypes (Fig. S4). We computed Pearson correlations among 8,971 genes for which there were non-zero counts in at least 10% of the cells from at least one cell type prior to imputation. We then applied k-means clustering (k=150) to the resulting gene co-expression matrix to derive gene modules. We dropped modules for which the first principal component (the module “eigengene”) explained less than 10% of the variance, and we merged modules whose eigengenes were >85% correlated. This resulted in a final set of 77 modules spanning 5,874 genes (Table S3).

Four analyses support the relevance of these gene co-expression modules to gene regulation and biology (Table S4). First, gene regulatory network reconstruction with GENIE3^31^ (using the smoothed expression profiles from the same cells and genes) revealed TF-target gene lists that significantly overlapped each of the 77 gene co-expression modules (FDR < 0.05), supporting the robustness of the modules and predicting specific TFs as key regulators of their activity. Second, all 77 modules also overlapped direct target genes of TFs inferred via motif analysis with RcisTarget^32^ (normalized enrichment score >= 3.71) and/or ChIP-seq data from ChEA^33^ (FDR < 0.05). Third, 75 of the 77 modules were enriched for at least one Gene Ontology functional annotation (p < 0.001). Fourth, all 77 modules overlapped a published gene co-expression module from bulk RNA-seq of striatal tissue in knock-in mouse models of the HD mutation^12^.

Notably, gene co-expression modules derived from single-nucleus RNA-seq appeared to have greater fidelity to specific cell types than the published network derived from bulk RNA-seq. For instance, genes from a large neuronally-enriched bulk RNA-seq gene co-expression module, “bulk M2”, previously shown to be down-regulated in HD knock-in mice, were enriched in 13 distinct snRNA-seq modules (FDR < 0.05; snRNA-seq modules M12, M50, M29, M101, M95, M86, M33, M46, M10, M70, M1, M76, and M36), all of which were down-regulated in *Htt^Q175/+^* vs. *Htt^+/+^* mice but with varying specificity across MSN subtypes and in other striatal cells (Fig. 3). Similarly, a down-regulated non-neuronal bulk RNA-seq module, “bulk M11”, overlapped five distinct snRNA-seq modules expressed specifically in astrocytes (M11), oligodendrocytes (M13, M31), or endothelial cells (M43), and across all glial cell types (M32). Thus, network reconstruction from single-cell RNA-seq provides complementary information about celltype-specific gene regulation that is not readily apparent from standard RNA-seq.

**Figure 3.**
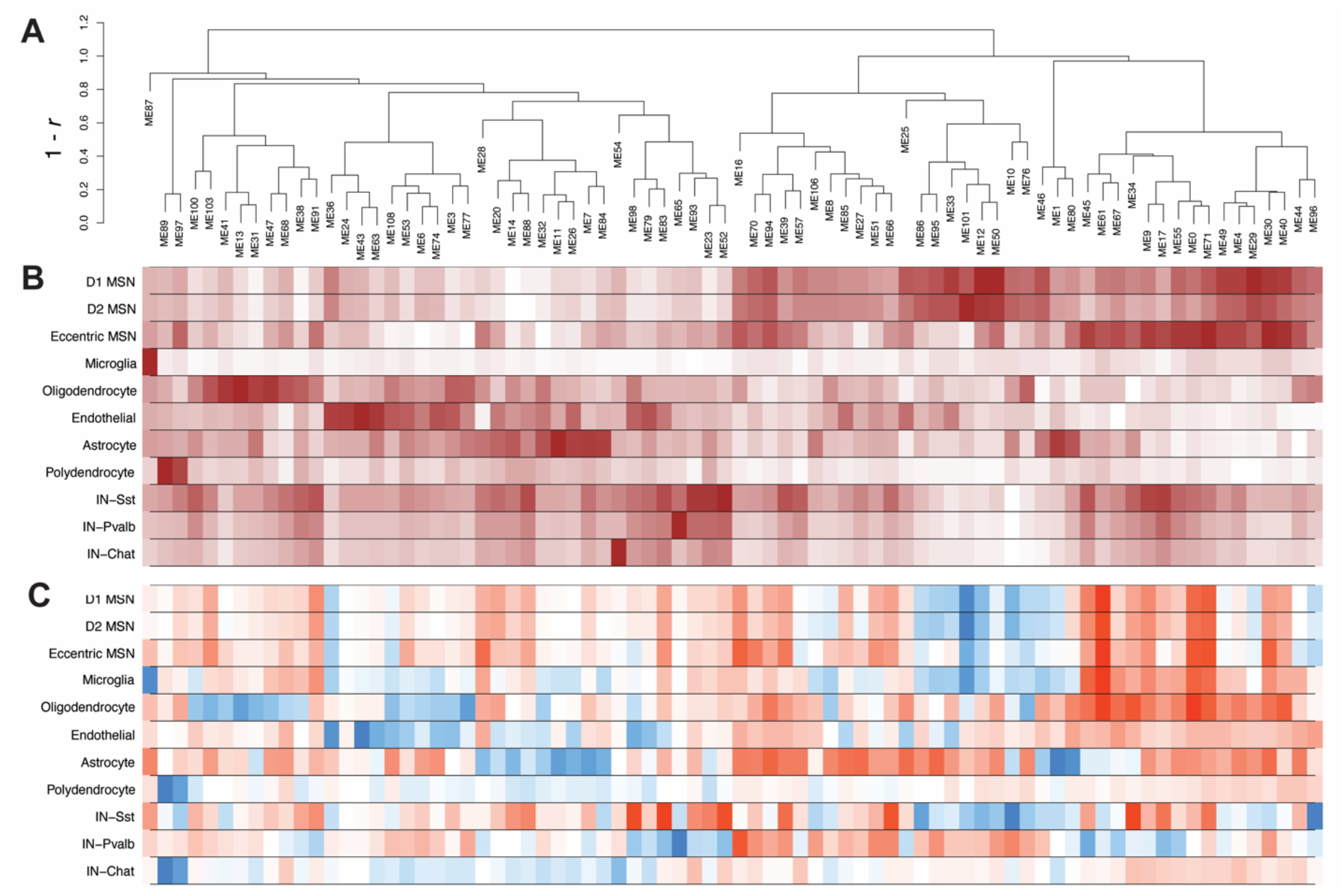
Identification of 77 gene co-expression modules in mouse striatum. A. Average-linkage hierarchical clustering of modules based on their first principal components (module eigengenes). B. Mean expression level of each module eigengine in each cell type. C. log2(fold change) of module eigengenes in cells from *Htt^Q175/+^* vs. *Htt^+/+^* mice.

We characterized the activity of the 77 gene co-expression modules across cell types and genotypes based on their eigengenes. 55 of the 77 modules were differentially expressed (adjusted p-value < 0.01, >1.5-fold change; Fig. 3c) in at least one cell type. These included 13 differentially expressed modules in D1 MSNs, 15 in D2 MSNs, 9 in eccentric MSNs, 39 in astrocytes, and 27 oligodendrocytes. Consistent with findings from DEGs (above), we identified numerous “cell identity” modules that were specifically expressed in one cell type and down-regulated in cells of that same type from *Htt^Q175/+^* vs. *Htt^+/+^* mice (Fig. 4, columns 1-4). We also identified modules that were broadly expressed and up-regulated across most or all cell types (Fig. 4, columns 5-6). A striking and unexpected finding was that many of the cell identity modules were also up-regulated in incorrect cell types. For instance, the MSN identity module M12 (Fig. 4, 1^st^ column) was down-regulated in D1 (logFC = −0.74, p < 1e-308), D2 (logFC = −0.74, p = 7.0e-217) and eccentric MSNs (logFC = −0.34, p = 3.7e-6), but up-regulated in astrocytes (logFC = 0.56, p-value = 3.5e-7) and oligodendrocytes (logFC = 0.27, p-value = 7.4e-3). The astrocyte identity module M11 (Fig. 4, 2^nd^ column) was down-regulated in astrocytes (logFC = −1.06, p = 2.1e-11), but slightly up-regulated in MSNs (logFC = 0.02, 0.03; p = 4.5e=10, 3.0e-4, in D1 and D2 subtypes, respectively). The oligodendrocyte identity module M13 (Fig. 4, 3^rd^ column) was down-regulated in oligodendrocytes (logFC = −0.96, p = 1.3e-9), but up-regulated in MSNs (logFC = 0.083, 0.08; p = 1.4e-27, 5.5e-18 in D1 and D2 subtypes) and astrocytes (logFC = 0.26, p = 2.0e-4). Further examples include the parvalbumin interneuron identity module M65 (Fig. 4; 4^th^ column) and the endothelial cell identity module M43, among others. Examining the expression of individual genes from these modules confirmed that they follow these same bi-directional patterns of transcriptional dysregulation (Fig. 4d). Notably, our gene regulatory network model predicted that many of these cell identity modules are regulated by canonical cell type-specific hub transcription factors, such as FOXP1 in M12, SOX9 in M11, MYRF in M13, and NKX2.1 (Fig. 4a,c), which are required for the development of MSNs, astrocytes, oligodendrocytes, and interneurons, respectively^34–37^.

**Figure 4.**
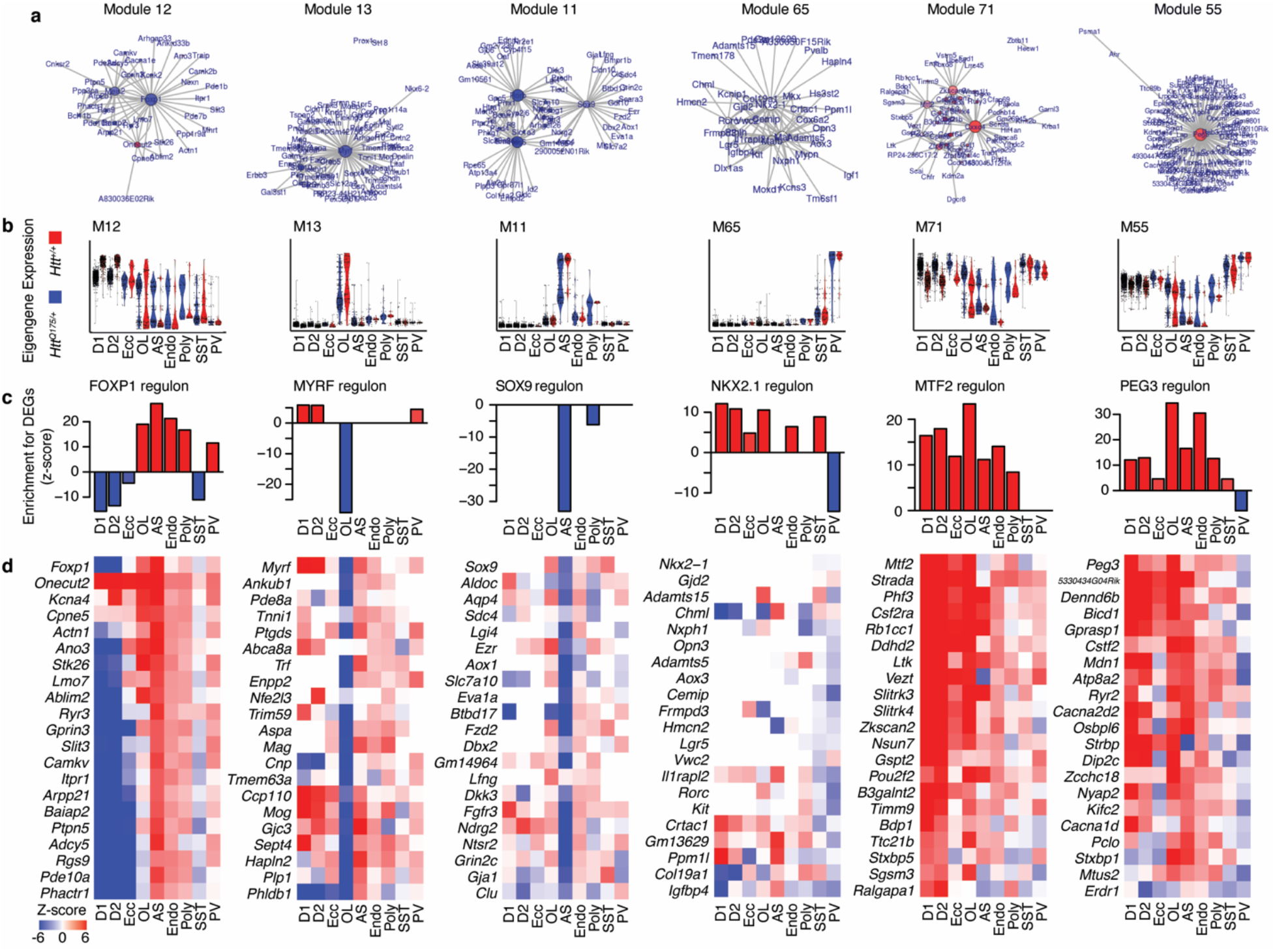
mHTT-associated cell type identity-related gene co-expression modules. a. Graphs of the top 100 gene-gene interactions within each module. Node size corresponds to eigencentrality. Node color corresponds to the log2(fold change) in expression between cells from *Htt^Q175/+^* vs. *Htt^+/+^* mice in the major cell type for that cluster (blue = down-regulated; red = up-regulated). b. Violin plots of module eigengenes. c. Over-representation of the predicted target genes from GENIE3 for the hub transcription factors in each module for up- and down-regulated genes in each cell type. d. Z-scores for cell type-specific up- or downregulation of the top 20 genes in each regulon shown in panel c.

We postulated that these bidirectional changes in gene expression may reflect aberrant repression and de-repression of cell type identity genes in *Htt^Q175/+^* mice. One mechanism by which this could occur is through interactions of wildtype and mutant HTT with Polycomb Repressive Complex 2 (PRC2). PRC2 facilitates gene repression via trimethylation of histone H3 at lysine 27 (H3K27me3), particularly in the promoters of genes involved in the development and maintenance of cell types. The HTT protein has genotype-specific interactions with PRC2 *in vitro^15^* and *in vivo*^38^, and conditional knockout of PRC2 in striatal MSNs causes gene expression changes that mimic the effects of HD mutations^39^. To test whether dysregulated cell type identity modules in *Htt^Q175/+^* mice involve PRC2, we assembled nine ChIP-seq datasets profiling the genomic occupancy for components of the PRC2 complex (EZH2, SUZ12) or for H3K27me3 in four disease-relevant cell types: medium spiny neurons^38,39^, astrocytes^40^, oligodendroocytes^27^, and embryonic stem cells^40,41^. We tested for over-representation of each of our 77 gene co-expression modules among putative PRC2 target genes, defined by the presence of a PRC2-related ChIP-seq peak +/- 5kb from a gene’s transcription start site. Ten modules were robustly over-represented for these PRC2 target genes (adjusted p-values < 0.01 in at least four of the ChIP-seq datasets; Fig. 5A). All of these ten PRC2-regulated modules were expressed specifically in a single striatal cell type (Fig. 5a), including modules specific to MSNs (M12, M29, M50), all interneurons (M23, M52), *Pvalb+* interneurons (M65), *Chat+* interneurons (M54), endothelial cells (M43), oligodendrocytes (M13), and astrocytes (M11). All of these PRC2-regulated modules except for those specific to interneurons were significantly down-regulated in that same cell type in *Htt^Q175/+^* mice (adjusted p-value < 0.01), while the expression of interneuron-specific modules trended downward in cells from *Htt^Q175/+^* mice (Fig. 5b). All ten PRC2-regulated modules (including interneuron-specific modules) were significantly up-regulated in at least one other cell type in which these genes are not normally expressed. As expected, the dynamics of PRC2 occupancy across cell types was negatively correlated with cell type-specific gene expression (Fig. 5c). PRC2 target genes in embryonic stem cells – in which genes for all differentiated cell types are repressed -- were over-represented in all ten modules. PRC2 target genes in MSNs were over-represented in interneuron- and glial-specific modules, but not in MSN-specific modules. PRC2 target genes in astrocytes and oligodendrocytes were primarily enriched in neuron-specific modules, but not in glial-specific modules.

**Figure 5.**
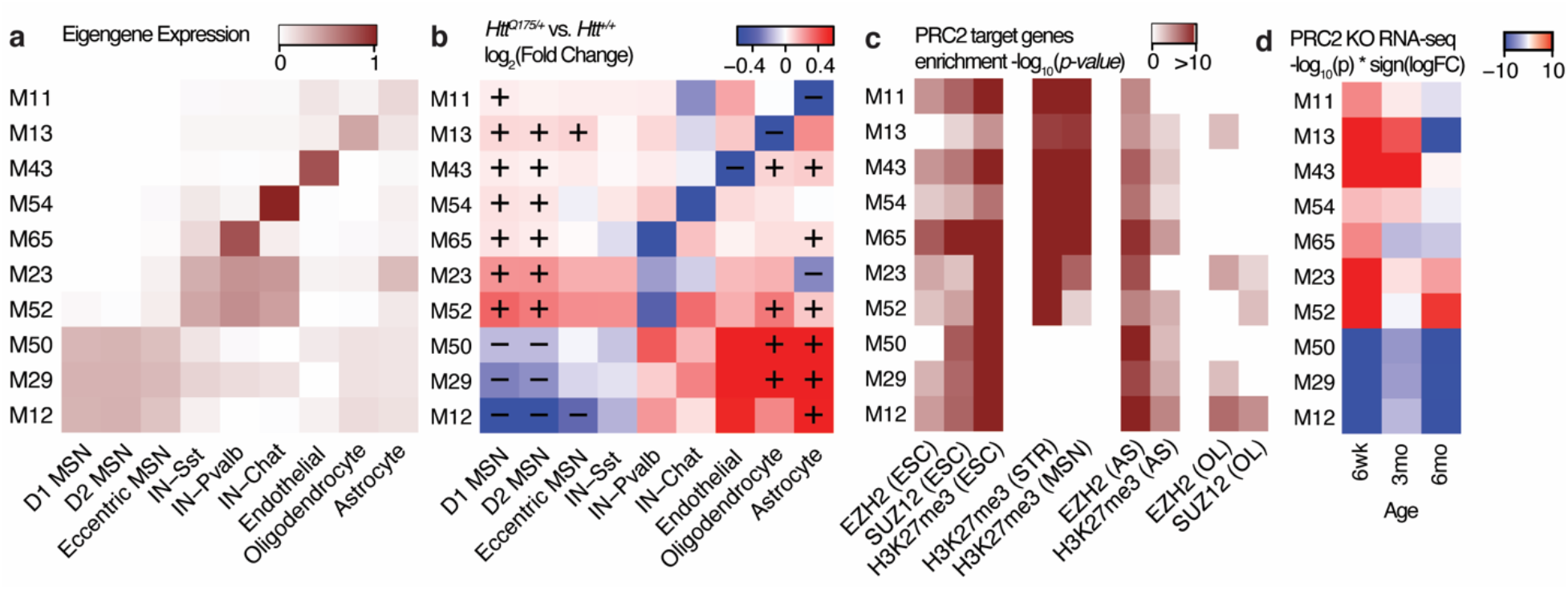
Regulation of HD-associated cell type identity modules by the Polycomb Repressive Complex 2 (PRC2). a. Cell type-specific expression of module eigengenes for ten gene co-expression modules robustly enriched for PRC2 target genes. b. log2(fold change) of module eigengenes in cells from *Htt^Q175/+^* vs. *Htt^+/+^* mice; “+” and “-” indicate statistically significant up- or down-regulation of eigengene expression in each cell types (adjusted p-value < 0.01. c. Over-representation of the genes in each module for PRC2 target genes defined by nine ChIP-seq experiments in embryonic stem cells (ESCs), MSNs, astrocytes (AS), and oligodendrocytes (OL). d. Over-representation of the genes in each module for up- and down-regulated genes in MSNs from 6-week-old, 3-month-old, and 6-month-old PRC2-silenced vs. control mice.

To more directly establish the regulation of these modules by PRC2, we analyzed published RNA-seq of MSNs from 6-week-old, 3-month-old, and 6-month-old EZH2/EZH1 double conditional knockout mice, in which PRC2 was silenced specifically in adult MSNs^39^. PRC2 silencing in MSNs strongly dysregulated all ten modules in the same direction observed in MSNs from *Htt^Q175/+^* vs. *Htt^+/+^* mice; i.e., modules specific to other cell types were ectopically expressed in PRC2-silenced MSNs, while MSN-specific modules were repressed (Fig. 5D). These analyses extend prior analyses of these data, which had also noted the bidirectional overlap with genes dysregulated in HD -- our analysis indicates that a much larger proportion of the transcriptional dysregulation in HD knock-in mice may be explained by altered interactions with PRC2 than had previously been appreciated and suggests that these interactions occur in multiple striatal cell types, not just MSNs. Thus, PRC2 regulates the cell type-specific expression patterns of many gene co-expression modules that are bidirectionally dysregulated in *Htt^Q175/+^* vs. *Htt^+/+^* mice, and our data support a model in which PRC2 loss-of-function due to the HD mutation causes both the derepression of these modules in inappropriate cell types, as well as their repression in their primary cell type.

## DISCUSSION

Here, we have described a comprehensive analysis of single-nucleus RNA-seq of striatal cells from a genetically precise knock-in mouse model of the HD mutation. Several novel findings merit particular attention. First, we observe shifts in cellular abundance revealing selective loss of D2 MSNs and increased microglia in aged *Htt^Q175/+^* mice. Second, we observe pronounced transcriptional dysregulation in nearly all striatal cell types, which we have here defined and compared to genes identified with RNA-seq in bulk striatal tissue. Third, analysis of HD-associated transcriptional networks across cell types reveals a striking pattern of loss of cell identity genes, coupled with aberrant up-regulation of these genes in incorrect cell types. Integration of these analyses with existing transcriptomic and epigenomic data suggest that altered PRC2 function may underlie these bidirectional changes, leading to dysregulation of cell identity across essentially all cell types in the adult striatum.

The early phases of disease progression in HD are dominated by hyperkinetic motor features, notably chorea. This has been attributed to the selective atrophy and early loss of D2 MSNs, revealed by the preferential loss of enkephalin-containing fibers across disease severity^42–44^. Our data support the higher vulnerability of this cell type, as we observed a selective reduced proportion of D2 MSNs. Despite the increased vulnerability of D2 MSNs, our data do not reveal significant differences among MSN subtypes with respect to the genes that are differentially expressed nor in the magnitude of the fold changes. The lack of quantifiable differences suggests that these responses to the mutation may be universal across all MSNs. However, we cannot rule out differences in the transcriptional responses of MSN subtypes at earlier time points, which should be examined in future studies.

The role of various types of glial cells in the progression of HD has been an important avenue of research. Our data reveal novel aspects of transcriptional dysregulation in glial cell types, which rely on the ability to examine transcriptional changes within specific cell types. Specifically, we find increased numbers of microglia, consistent with previous reports that microgliosis begins in presymptomatic HD mutation carriers^6^. In addition, we detected thousands of celltype-specific DEGs in glia -- especially astrocytes and oligodendroocytes -- many of which are not readily apparent in RNA-seq of bulk striatal tissue. Across glial cell types, we observed a spectrum of transcriptional states, comprising of nuclei in similar transcriptional states as the wild type nuclei, nuclei in discernably different transcriptional states as well as intermediate states between the two that may indicate a gradual shift in cell states and opens new areas of study. This is distinct from our trajectory analysis of MSNs, in which we observed nearly complete separation of wildtype and *Htt^Q175/+^* cellular states. Activation of neuroinflammatory genes is observed across several cell types, including astrocytes, oligodendrocytes, and MSNs. However, the transcriptional states of glia are distinct from classical neuroinflammatory states such as ‘A1’ or ‘A2’ reactive astrocytes, suggesting disease-specific mechanisms.

Across cell types, a common feature of the transcriptional dysregulation we observe is aberrant maintenance of cell fate commitment. Previous studies, based on RNA-seq of bulk striatal tissue, have suggested that loss of cell fate commitment may drive MSN vulnerability in HD^12^. Reconstruction of gene co-expression networks and gene regulatory networks from our data revealed numerous gene modules that were dysregulated in HD knock-in mice. HD-associated gene modules that are expressed primarily in MSNs overlapped strongly with disease-associated gene modules from bulk RNA-seq of HD and HD mouse models, which is to be expected since MSNs represent a large percentage of cells in the striatum. However, the improved resolution of our network models across cell types provides greater insight into this mode of pathology. Notably, we confirm that aged MSNs expressing mHTT have reduced expression of MSN-specific genes, e.g. the FOXP1 regulon in module M12. In addition, our findings reveal reduced cellular identity genes in other cell types, including oligodendrocytes (MYRF regulon in module M13) and astrocytes (SOX9 regulon in module 11). Here we observe, for the first time, that in addition to being downregulated in their correct cell type, cell identity genes are also aberrantly up-regulated in other cell types. This expands the model of cell fate commitment changes in HD, suggesting it may be a shared feature of aged cells expressing mHTT, affecting virtually all cell types in the striatum.

Given the importance of PRC2 in establishing and maintaining cell fate and the known interactions between HTT and PRC2, we compared our results to PRC2-related ChIP-seq and RNA-seq datasets. We find that ten of our 77 gene co-expression modules are enriched with PRC2 target genes, all of which were expressed and down-regulated in specific cell types. Our results suggest that PRC2 helps restrict expression of these genes to their appropriate cell type in the adult striatum. Conditional knockout of PRC2 and the presence of mHTT each results in highly similar patterns of bidirectional dysregulation across cell types. Thus, many of the transcriptional effects of mHTT may be mediated by PRC2 loss-of-function. Consistent with this idea, we recently described a global decrease of H3K27me3 levels in striatal tissue from HD knock-in mice^38^. Future studies should test whether restoring PRC2 function can rescue the phenotypic consequences of HD mutations.

Our study has several limitations. The relatively small size of our dataset limited our ability to characterize gene expression changes in rare cell types, including microglia. In addition, our analysis is limited to a single mouse model at a single time point in a single tissue. There will be value in generating additional single-nucleus data, including from earlier time points and other disease-affected brain regions, and to expand these analyses to human cells and brain regions. In summary, our study demonstrates the power of single-nucleus RNA-seq to reveal novel insights into celltype-specific mechanisms in HD, many of which were unknown despite decades of studies on the HD mutation.

## METHODS

### Mice

*Htt^Q175/+^* mice and wildtype littermate controls on the C57BL6/J background (B6J.129S1-*Htt^tm1.1Mfc^*/190ChdiJ; JAX stock #029928) were bred and aged in the colony of the Carroll lab at Western Washington University. The B6J.zQ175DN KI used in this study also lack a floxed neo cassette 1.3 kbp upstream of exon 1 that is not known to have an effect on the phenotypic characteristics of the mice. Three 14-month-old and one 15-month-old male mice of each genotype were used to generate the primary dataset described here. Mice were deeply anesthetized using a phenobarbital based euthanasia solution (Fatal Plus, Henry Schein) and striatal tissue dissected on ice. Tissue was immediately flash frozen in liquid nitrogen and stored at −80°C. Experiments were performed following NIH animal care guidelines and approved by Western Washington University’s Institutional Animal Care and Use Committee under protocol 16-011.

### Isolation of nuclei from frozen brain tissue

Nuclei were isolated from flash-frozen striatal tissue as described in previous protocols^45–47^, with slight modifications. A detergent-mechanical cell lysis method was used, involving 3 major steps: lysis, homogenization and density barrier centrifugation. In a laminar hood, a ~50 mg piece of frozen brain tissue was placed into a pre-frozen BioPulverizer (BioSpec) and smashed to a thin frozen layer, then immediately transferred to lysis buffer (250mM Sucrose, 25mM KCl, 5mM MgCl2, 1uM DTT, 1X RNAse Inhibitor, 0.1% TritonX-100, 10mM Tricine Buffer, pH 8.0). Tissue was disaggregated, flushing up and down, first using a 1 ml pipette tip, then with a 30G needle in a 3ml syringe. Homogenized tissue was diluted to 10ml with lysis buffer and filtered through a 70 micron filter. Homogenate was spun at 1,000g for 8 min at 4°C. Pellet was then re-suspended and 500 μl of pellet suspension was diluted with 500 μl 50% Iodixanol Solution (50% Iodixanol, 250mM Sucrose, 150 mM KCl, 30mM MgCl2, 1X RNAse Inhibitor, 60mM Tricine Buffer, pH 8.0), filtered for second time with a 70 micron mesh and placed on top of 500 μl layer of 29% Iodixanol Solution (29% Iodixanol, 250mM Sucrose, 150 mM KCl, 30mM MgCl2, 1X RNAse Inhibitor, 60mM Tricine Buffer, pH 8.0). Density barrier was centrifuged at 13,500g for 20 min at 4°C. The pellet was collected and washed with 10 ml of PBS, 2% BSA, 1X RNAse Inhibitor, centrifuged at 1,000g for 8 min at 4°C, and finally suspended in 1 ml of PBS, 2% BSA, 1X RNAse Inhibitor and filtered using a 40 micron Flowmi cell strainer. Single nuclei suspensions were counted and evaluated for integrity using Propidium Iodide in a MoxiGo cytometer using 650 nm filter. Nuclei count was adjusted to 5000 nuclei/ml.

### Library preparation and sequencing

13,000 nuclei were loaded into each well of a Chromium microfluidics controller (10X Genomics, Pleasanton, CA) using PBS + 2% BSA. Sequencing libraries were generated using the Chromium Single Cell Gene Expression 3’ kit with Version 3 chemistries. Samples were sequenced across two lanes of an Illumina HiSeq4000 sequencer to obtain 75 base pair paired-end reads.

### Cell QC and data pre-processing

Raw sequencing reads were processed to counts of unique molecular identifiers (UMIs) in each droplet with cellranger v3.0.2 (10X Genomics). Artifacts from ambient RNA were reduced with SoupX^48^, based on a uniform contamination fraction estimate of 10%. This contamination estimate was derived using marker genes >20-fold enriched in each cell type from DropViz^25^. Also using SoupX, we derived the following sample-specific thresholds at which we observed relatively low contamination, and we removed cells outside these ranges: 1260-6300 UMIs/cell for BBY2, BCR2, BCR3, BCR4, BCR5 and BCR6, and 1260-4470 nUMIs for BCR1. One *Htt^+/+^* sample, BBY3, was deemed to be an outlier based on very low UMI counts across most cells and was dropped from further analysis. In addition, using Seurat, we removed libraries with >5% of read counts from mitochondrial genes, as the presence of these non-nuclear transcripts indicates both incomplete fractionation of nuclei and cellular stress. To obtain the final set of cells for clustering, we manually removed any remaining doublets with high expression of mutually exclusive markers.

### Normalization and batch effect correction

UMI counts from each cell were normalized to log (counts per million). A uniform set of highly-variable genes was selected by identifying the top 5000 most variable genes in each sample separately using the FindVariableFeatures() function in Seurat, then taking the intersect across samples, resulting in a final set of 937 highly-variable genes. Expression levels for these highly-variable genes were centered and scaled within each sample. Then, batch effects correction was performed across samples with MNN^49^, implemented with the mnnCorrect() function in the scran R package. MNN normalization is performed sequentially across samples and lacks a built-in function to optimize sample order. Therefore, 17 MNN iterations were performed using different sample ordering, and the best ordering was selected based on cell clustering results. In the final ordering, samples were in the order BBY2, BCR1, BCR2, BCR3, BCR4, BCR5 and BCR6.

### Cell type clustering and labeling

MNN-adjusted counts from 5,429 cells passing the initial QC were used for clustering, using the RunPCA(), FindNeighbors(), and FindClusters() functions in Seurat v3.0. The clusters were visualized using the RunTSNE() function in Seurat. npcs=10 principal components, k=30 shared nearest neighbors, and a Louvain clustering resolution of 0.4 were empirically determined to produce optimal clustering. Marker genes identified in each cluster using the FindAllMarkers() function were then used to assign a cell type label to each molecularly-defined cluster based on known marker genes for striatal cell types (dropviz.org). Three clusters were dropped from further analysis because they included cells from only one sample, were marked by high expression of mitochondrial transcripts, or expressed markers from mixed cell types, resulting in the final set of 4,524 cells.

Identifying subtypes of medium spiny neurons presented a particular challenge, since the small number of markers distinguishing D1 vs. D2 MSNs were dwarved by the massive effects of HD mutations in these cells. Therefore, we performed sub-clustering of MSNs using 36 marker genes with >8-fold difference in expression between D1 vs. D2 MSNs in the DropViz atlas^25^. Using the normalized counts from these 36 genes, we computed pairwise Spearman rank correlations among all canonical MSNs, and we performed average-distance hierarchical clustering using 1 – Spearman’s rho as a distance metric, revealing two major groups of cells with high expression of markers for D1 vs. D2 MSNs, respectively. 87 MSNs were in smaller clusters that lacked high expression for either subtype and were discarded.

Similarly, clusters of reactive vs. non-reactive astrocytes were derived by sub-clustering with 87 markers of reactive and non-reactive subtypes^28^. In this case, sub-clustering was performed in Seurat v3, using the top 5 principal components and a Louvain modularity resolution of 0.5.

### Smoothing of read counts

For the purpose of assessing gene expression differences between genotypes, and subsequent analyses of pseudotime trajectories and gene co-expression clustering, we smoothed read counts to reduce dropout effects and improve gene-gene correlation structure. Smoothing was performed with knn-smoothing^30^, using k=15 neighbors and 30 PCs. Smoothed counts were normalized using the Seurat NormalizeData() function for downstream analyses.

### Celltype-specific differentially expressed genes

We identified celltype-specific differentially expressed genes (DEGs) in *Htt^Q175/+^* vs. wildtype mice by three methods. In our primary analysis, we compared cells of each genotype using Wilcoxon Rank Sum tests implemented with the Seurat FindMarkers() function, testing all genes with non-zero imputed counts in at least 10% of the cells from at least one of the two genotypes. This analysis treats the cell (not the mouse) as the primary unit of analysis. We believe this choice is justified, as both the neurodegenerative processes in MSNs and the activation of inflammatory states in glial cells are thought to occur in a relatively cell-autonomous fashion, resulting in a mosaic of cell states within each mouse that would not be captured in an analysis treating the mouse as the primary unit of analysis. To confirm that DEGs detected by our primary approach were robust, we also conducted a secondary analysis in which the Seurat FindMarkers() function was applied to non-smoothed read counts.

### Trajectory analysis

We used Seurat v3 to perform principal components analysis on the centered and scaled expression levels of the 5,000 most variable genes within each cell type. We eliminated any principal component for which the strongest loadings were dominanted by mitochondrial genes, as these vectors typically correspond to technical variation among cells rather than biological signal. Using the embeddings of the top five remaining principal components, we then identified a non-branching pseudotime trajectory with the slingshot() function in the Slingshot v1.4.0 R package^26^. We tested for associations of slingPseudoTime and genotype using the t.test() function in R.

### Gene set enrichment analysis

DEGs in each cell type and genes in each gene co-expression module were tested for enrichment in curated gene sets from four sources. First, we tested for enrichments in 2,368 curated gene sets from the Huntington’s Disease molecular Signatures Database (https://www.hdinhd.org/2018/05/22/hdsigdb/), including lists of DEGs and WGCNA modules from RNA-seq experiments, as well as known marker genes for striatal cell types. Second, we tested for enrichments in 12,177 Gene Ontology terms from the org.Mm.egGO2ALLEGS object in the org.Mm.eg.db R package. Third, we tested for enrichments in transcription factor regulons from ChIP-seq experiments in ChEA^33^ (https://amp.pharm.mssm.edu/Enrichr/geneSetLibrary?mode=text&libraryName=ChEA_2016). Fourth, we tested for enrichments with evolutionarily conserved sequence motifs from RcisTarget^32^. For the first three sources, enrichments among down- and up-regulated genes were tested separately using the geneSetTest() function in the limma R package with ranks.only = TRUE and type = “t” and using genes ranked from most strongly down-regulated to most strongly up-regulated based on the −log10(p-value) multiplied by the sign of the log2(fold change). Motif analysis was performed with RcisTarget using default parameters.

### Statistical overlap of celltype-specific differentially expressed genes across cell types

Statistical overlap among ranked lists of down- and up-regulated genes in each pair of cell types was evaluated using rank-rank hypergeometric overlap (RRHO), implemented using the RRHO() function from the RRHO R package^50^. The RRHO algorithm steps through two gene lists ranked by the degree of differential expression in two independent experiments, successively measuring the statistical significance of the number of overlapping genes. Each comparisoin between cell types used the set of genes expressed with a non-zero read count in at least ten percent of the cells form both cell types, with the genes ranked from most strongly down-regualted to most strongly up-regulated within each cell type based on the −log10(p-value) multiplied by the sign of the log2(fold change). We used a step size such that each gene list was divided into 100 equally sized bins.

### Gene co-expression modules

Gene-gene correlation structure in single-nucleus RNA-seq is notoriously weak due to the sparsity of the data. Therefore, for network analyses we used smoothed counts, as described above, to impute a more complete representation of gene expression in each cell. We selected 8,971 genes with non-zero counts in at least 10% of the cells from at least one cell type. We computed Pearson correlations among all pairs of genes. We then applied k-means clustering to the resulting gene coexpression matrix to derive gene modules. k=150 was manually determined to capture some of the finer structure in the network without over-splitting. Each module’s eigengene -- the first principal component - was computed with the moduleEigengenes() function in the WGCNA R package. We dropped modules for which the eigengene explained less than 10% of the variance, and we merged modules whose eigengenes were >85% correlated. These procedures resulted in a final set of 77 modules spanning 5,874 genes.

### Gene regulatory network

A gene regulatory network model describing predicted interactions between TFs and their potential target genes in the mouse striatum was derived using GENIE3^31^. The GENIE3 algorithm performs network reconstruction by using a random forest regression model to select sets of TFs whose combined expression predicts the expression of each gene. We started with the smoothed counts of 8,971 genes with non-zero counts in at least 10% of the cells from at least one cell type, as for gene coexpression networks above. We downloaded a curated list of all 1639 known and likely human TFs from http://humantfs.ccbr.utoronto.ca/download/v_1.01/TFs_Ensembl_v_1.01.txt. We identified 1,373 mouse orthologs of these TFs using the biomaRt R package (accessed April 21, 2020). We then intersected this list of putative mouse TFs with the 8,971 striatally-expressed genes from our dataset, producing a list of of 589 striatally-expressed TFs. GENIE3 was run using default parameters as implemented in the GENIE3 BioConductor package v1.1. The algorithm produces a very long list of potential TF-gene interactions ranked by the randomForest importance score, and many of these potential interactions are very weak. This importance score is agnostic to whether the TF-gene interaction is positive (“activating”) or negative (“inhibitory”), but it has been suggested that inhibitory interactions are less reliable. We therefore retained only the predicted interactions between pairs of genes whose expression were positively correlated (Pearson’s r > 0), and we trimmed this list to the top remaining 180,000 TF-gene interactions with the strongest importance scores, corresponding to a mean in-degree of ~20 TFs per gene.

### Overlap with Polycomb Repressive Complex 2 (PRC2)-related datasets

We analyzed PRC2 target genes derived from nine ChIP-seq datasets in four cell types to test for over-representation in gene coexpression modules. When available, we used published target gene lists. Otherwise, we downloaded aligned sequence reads, performed peak-calling with MACS v2.1^51^, and annotated peaks to genes with transcription start sites within +/- 5 kb. The nine datasets are as follows: (i) ChIP-seq of EZH2 in mouse embryonic stem cells^41^, obtained from ChEA^33^; (ii) ChIP-seq of SUZ12 in mouse embryonic stem cells^41^, obtained from ChEA^33^; (iii) ChIP-seq of H3K27me3 in mouse embryonic stem cells, generated by Bing Ren’s lab (UCSD) for the ENCODE consortium (ENCFF055QNY); (iv) ChIP-seq of H3K27me3 in mouse MSNs^39^, obtained from HDSigDB; (v) ChIP-seq of H3K27me3 in bulk striatal tissue from four-month-old *Htt^Q111/+^* and *Htt^+/+^* mice^38^; (vi) ChIP-seq of EZH2 in human astrocytes, generated by Bradley Bernstein’s lab for the ENCODE consortium (reproducible peaks from ENCFF254DFD and ENCFF831JFC); (vii) ChIP-seq of H3K27me3 in human astrocytes, generated by Bradley Bernstein’s lab for the ENCODE consortium (ENCFF315BVX); (viii) ChIP-seq of EZH2 in mouse corpus callosum (enriched for oligodendrocytes; Table S3 from ^27^); (ix) ChIP-seq of EZH2 in mouse corpus callosum (enriched for oligodendrocytes; Table S3 from ^27^). We tested for over-representation in gene co-expression modules using Fisher’s exact tests. We also analyzed published microarray gene expression profiles (Affymetrix 430_2 array) of MSNs from 6-week-old, 3-month-old, and 6-month-old *Ezh1^-/-^; Ezh2^fl/fl^; Camk2a-cre* vs. control mice, in which PRC2 had been conditionally silenced in adult MSNs^39^. Normalized data were downloaded from the Gene Expression Omnibus (GSE84243). We fit a linear model using the lmFit() function in the limma R package, followed by post-hoc contrasts to estimate the effect of genotype at each time point, using contrasts.fit() and eBayes(). Over-representation analysis of gene co-expression modules among up- and down-regulated genes was performed with the geneSetTest() function.

## AUTHOR CONTRIBUTIONS

S.A.A. and J.B.C. designed the study; M.C.-G. performed molecular and cellular experiments; S.R.C., S.R.W.L., and J.P.C. performed mouse work; S.M., B.R.H., C.C., and S.A.A. analyzed the data; S.M., S.A.A. and J.B.C. drafted the manuscript; all authors reviewed the manuscript.

## DATA AVAILABILITY

Raw and processed data have been submitted to the Gene Expression Omnibus (accession # in progress). A web portal for visualization and analysis of these data is available via the Gene Expression Analysis Resource (https://umgear.org/p?l=1d76bf3e).

## ACKNOWLEDGEMENTS

This work was supported by contracts from the CHDI Foundation (S.A.A. and J.B.C., PIs) and by a grant from the BRAIN Initiative (R24 MH114815, Ronna Hertzano, PI).

## SUPPLEMENTARY INFORMATION

**Figure S1.**
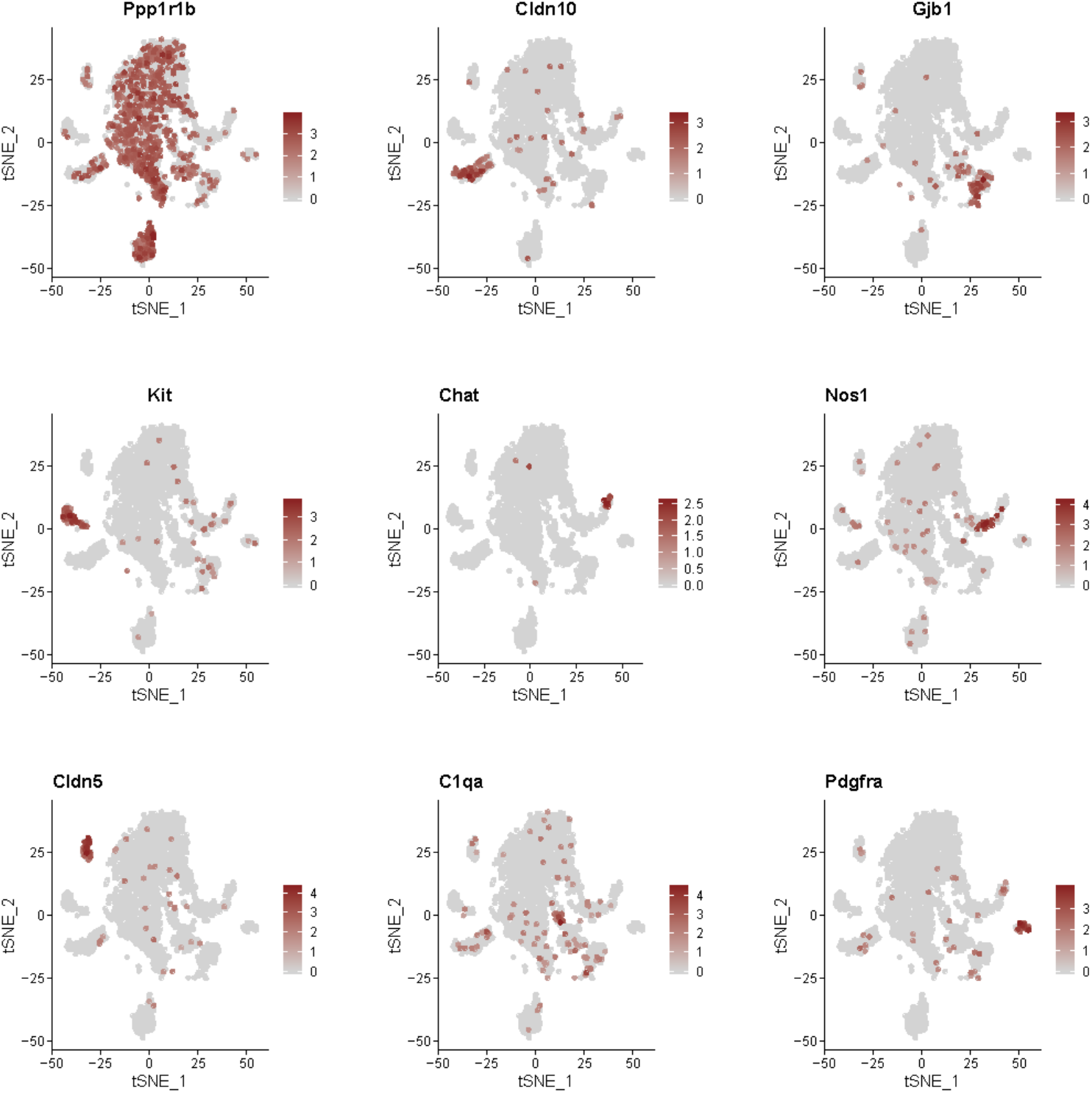
Expression of cell type marker genes. From left to right and top to bottom: *Ppp1r1b* (medium spiny neurons); *Cldn10* (astrocytes); *Gjb1* (oligodendrocytes); *Kit (Pvalb+* interneurons); *Chat (Chat+* interneurons); *Nos1* (*Sst+* interneurons); *Cldn5* (endothelial cells); *C1qa* (microglia); *Pdgfra* (polydendrocytes).

**Figure S2.**
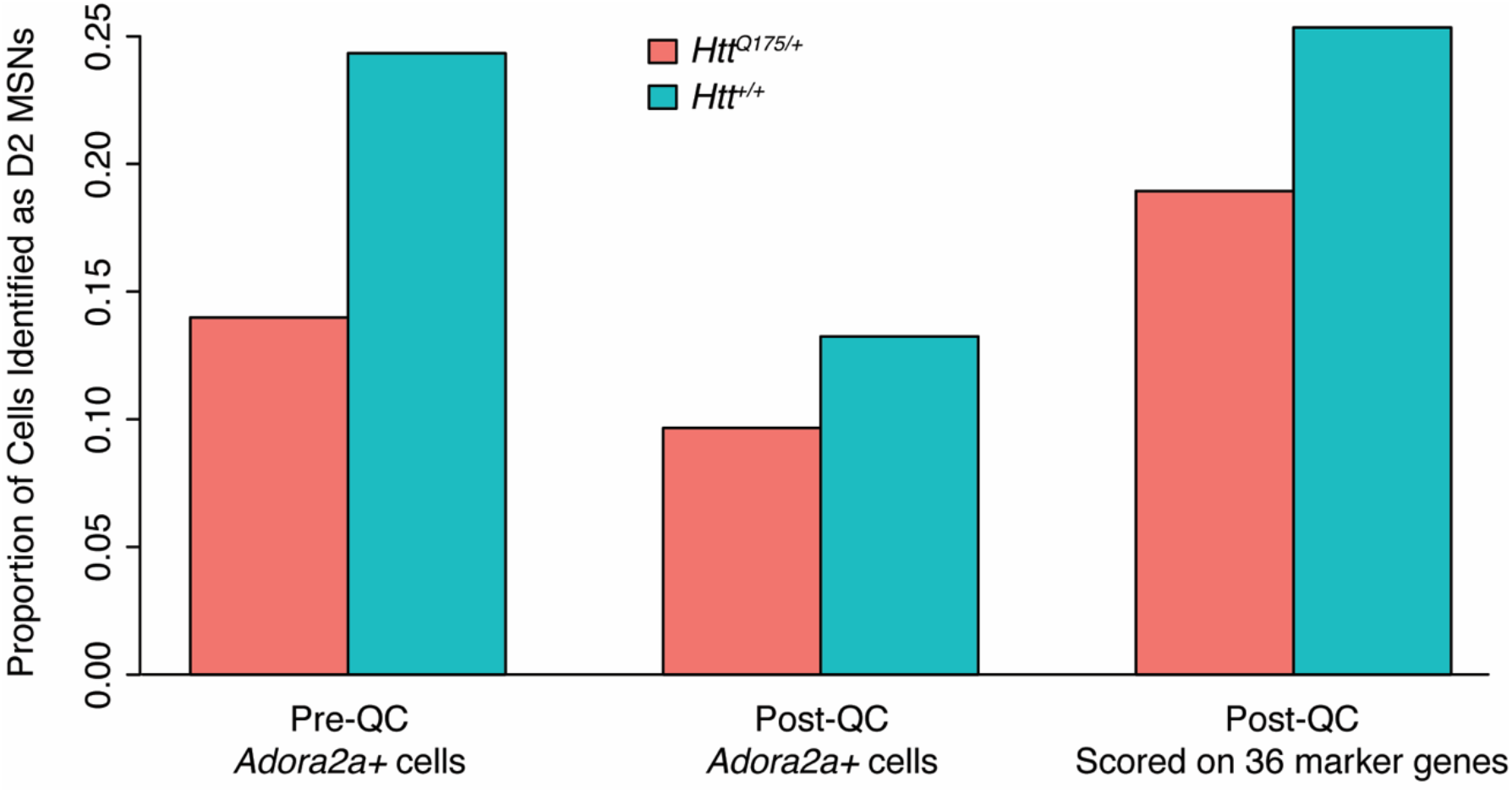
Decreased proportion of D2 MSNs in *Htt^Q175/+^* vs. *Htt^+/+^* mice is robust to parameters. Left: Proportion of cells identified as D2 MSNs in raw data (prior to QC) based on expression of *Adora2a*. Middle: Proportion of cells identified as D2 MSNs in QC-ed based on expression of *Adora2a*. Right: Proportion of cells identified as D2 MSNs in QC-ed based on expression of 36 marker genes (duplicated from Fig. 1d).

**Figure S3.**
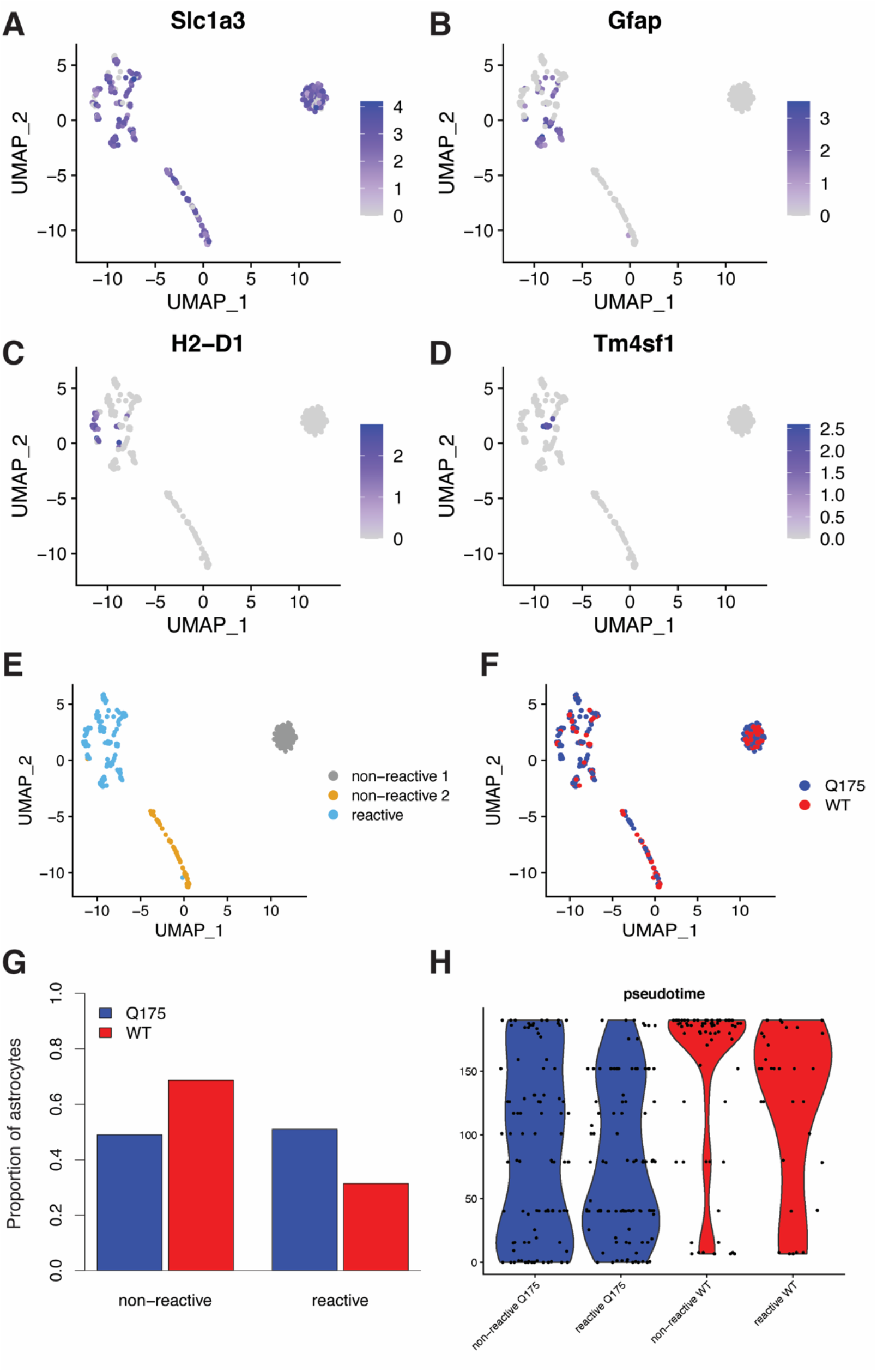
Characterization of reactive astrocytes in the striatum of *Htt^Q175/+^* vs. *Htt^+/+^* mice. Reactive astrocytes were identified by sub-clustering astrocytes on the expression of 87 markers of reactive populations. a-d. Expression of marker genes for all astrocytes (a, *Slc1a3*), pan-reactive astrocytes (b, *Gfap*), A1 reactive astrocytes (c, *H2-D1*), and A2 reactive astrocytes (d, *Tm4sf1*). e. Assignments of astrocyte sub-clusters as reactive vs. non-reactive based on known markers. f. Distribution of cells from *Htt^Q175/+^* vs. *Htt^+/+^*mice across astrocyte sub-clusters. g. Proportion of reactive vs. non-reactive astrocytes in *Htt^Q175/+^* vs. *Htt^+/+^*mice. h. Violin plots showing the relationship between the reactive state and genotype of astrocytes and Slingshot pseudotime.

**Figure S4.**
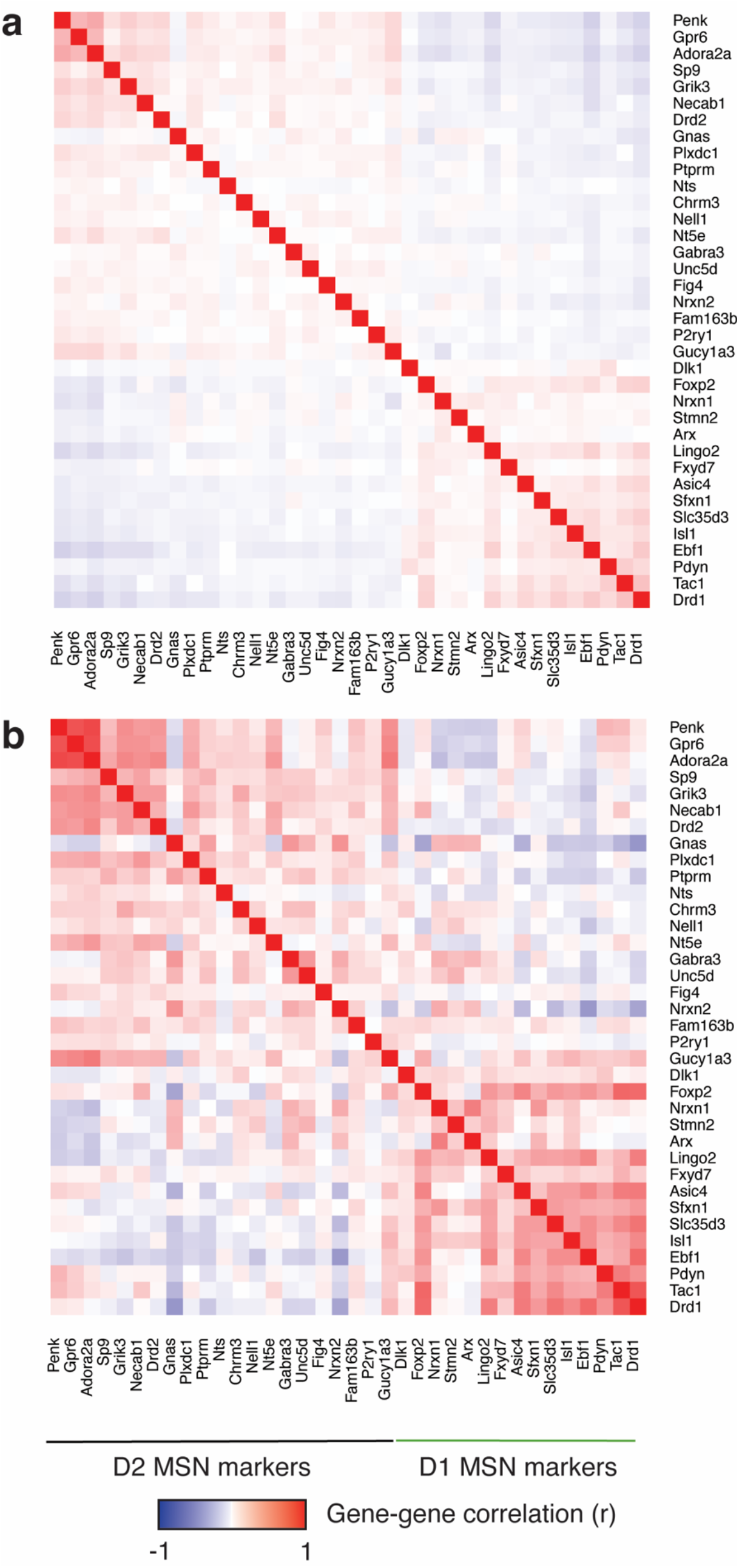
Smoothing rescues gene-gene correlation structure among markers of D1 vs. D2 MSNs. a. Pearson correlations among 36 marker genes for D1 vs. D2 MSNs in normalized, non-smoothed counts from the 4,524 cells in our dataset. b. Pearson correlations among the same 36 marker genes using smoothed counts. The 36 marker genes are the same genes shown in Fig. 1c and are ranked in order of subtype-specificity based on data from the DropViz atlas.

**Table S1. Cell type marker genes and differentially expressed genes in nine striatal cell types from 14-15-month-old *Htt^Q175/+^* vs. *Htt^+/+^* mice.** Tab 1. Marker genes in each cell type. Tab 2. DEGs in each cell type based on smoothed counts. Tab 3. DEGs in each cell type based on non-smoothed counts. (Excel file)

**Table S2. Gene set enrichment analysis of celltype-specific differentially expressed genes.** Enrichments in gene sets from Gene Ontology (tab 1) and HDSigDB (tab 2), and for the predicted target genes of each striatally expressed transcription factor in our GENIE3 gene regulatory network model (tab 3). (Excel file)

**Table S3. Gene membership in 77 gene co-expression modules**. (Excel file)

Table S4. **Gene set enrichment analysis of** of gene co-expression modules. Enrichments were calculated in the following gene sets: Tab 1. Gene Ontology terms. Tab 2. HDSigDB terms. Tab 3. WGCNA modules from Langfelder et al. 2016 (subset of HDSigDB). Tab 4. Predicted target genes of TFs based on ChIP-seq experiments in ChEA. Tab 5. Predicted target genes of striatally expressed TFs in our GENIE3 gene regulatory network model. Tab 6. Evolutionarily conserved cis-motifs from RcisTarget. (Excel file)

## Notes

### Competing Interest Statement

The authors have declared no competing interest.

